# Impact of a human gut microbe on *Vibrio cholerae* host colonization through biofilm enhancement

**DOI:** 10.1101/2021.02.01.429194

**Authors:** Kelsey Barrasso, Denise Chac, Meti D. Debela, Catherine Geigel, Jason B. Harris, Regina C. LaRocque, Firas S. Midani, Firdausi Qadri, Jing Yan, Ana A. Weil, Wai-Leung Ng

**Affiliations:** Department of Molecular Biology and Microbiology, Tufts University School of Medicine, Boston, MA; Program of Molecular Microbiology, Graduate School of Biomedical Sciences, Tufts University School of Medicine, Boston, MA; Department of Medicine, University of Washington, Seattle, WA; Department of Molecular, Cellular and Developmental Biology, Yale University, New Haven, CT; Quantitative Biology Institute, Yale University, New Haven, CT; Division of Infectious Diseases, Massachusetts General Hospital, Boston, MA; Department of Pediatrics, Harvard Medical School, Boston, MA; Department of Molecular Virology and Microbiology at Baylor College of Medicine, Houston, TX; International Center for Diarrheal Disease Research, Bangladesh, Dhaka, Bangladesh

**Keywords:** Bacterial pathogenesis, gut microbiome, mixed species biofilm

## Abstract

Recent studies indicate that the human intestinal microbiota could impact the outcome of infection by *Vibrio cholerae*, the etiological agent of the diarrheal disease cholera. A commensal bacterium, *Paracoccus aminovorans*, was previously identified in high abundance in stool collected from individuals infected with *V. cholerae* when compared to stool from uninfected persons. However, if and how *P. aminovorans* interacts with *V. cholerae* has not been experimentally determined; moreover, whether any association between this bacterium alters the behaviors of *V. cholerae* to affect the disease outcome is unclear. Here we show that *P. aminovorans* and *V. cholerae* together form dual-species biofilm structures at the air-liquid interface, with previously uncharacterized novel features. Importantly, the presence of *P. aminovorans* within the murine small intestine enhances *V. cholerae* colonization in the same niche that is dependent on the *Vibrio* exopolysaccharide (VPS) and other major components of mature *V. cholerae* biofilm. These studies illustrate that dual-species biofilm formation is a plausible mechanism used by a gut microbe to increase the virulence of the pathogen, and this interaction may alter outcomes in enteric infections.

**Significance Statement:** While ample evidence suggests that the outcome of some enteric infections can be affected by the intestinal microbiota, how specific gut microbes change the behaviors of a pathogen is unclear. Here we characterize the interaction between *Vibrio cholerae* and *Paracoccus aminovorans*, a gut microbe known to increase in abundance in the intestines during active *V. cholerae* infection in humans. These two bacteria form a dual-species biofilm structure at the air-liquid interface, and the gut microbe increases the host colonization efficiency of *V. cholerae*. Importantly, our study identifies a previously unknown mechanism of gut microbe-pathogen interaction that has the potential to alter the disease outcome.

## Introduction

*Vibrio cholerae* (*Vc*) causes an estimated 3 million infections and 120,000 deaths each year, and larger and more deadly outbreaks have increased during the last decade (1, 2). A wide range of clinical outcomes occur in persons exposed to *Vc*, ranging from asymptomatic infection to severe secretory diarrhea. It is nearly certain that many behaviors of *Vc* in the aquatic environment and inside the host are significantly affected by the presence of other microbes (3), and recent studies provide evidence that the gut microbiota may impact the severity of cholera (4–7).

Several functions of the gut microbiota influence the growth or colonization of enteric pathogens, including production of anti-microbial compounds, maintenance of the intestinal barrier, regulation of the host immune response, and modulation of available nutrients (8). Gut microbes have been shown to have an important role in *Vc* infection in various animal models. For instance, disruption of the commensal microbiota with antibiotics is required to allow successful *Vc* colonization in adult rodent models (9, 10). Conversely, *Vc* actively employs a Type VI secretion system to attack host commensal microbiota to enhance colonization of the gut in infant mice (11). Moreover, specific microbial species have a profound impact on *Vc* colonization. *Blautia obeum*, an anaerobic Gram-positive bacterium, decreases *Vc* intestinal colonization presumably by producing a signaling molecule that induces *Vc* into a high cell-density quorum sensing state (5) in which virulence gene expression is repressed (12, 13). Certain microbiota species reduce *Vc* colonization by producing the enzyme bile salt hydrolase that degrades the host-produced virulence-activating compound taurocholate (4, 5). Through metabolizing host glycans into short chain fatty acids that suppresses *Vc* growth, a prominent commensal species, *Bacteroides vulgatus*, reduces *Vc* proliferation within the intestine (14).

While the above studies exemplify how a single microbe or a group of microbes can protect the host from *Vc* infection, the mechanisms used by certain gut microbes to promote *Vc* virulence, thereby increasing the likelihood of individuals to develop cholera and worsen disease outcomes, are less well understood. We have previously studied household contacts of cholera patients to understand how gut microbes impact on susceptibility to cholera and identified bacteria associated with increased or decreased susceptibility to *Vc* infection (6, 7). We also observed that the gut microbial species *Paracoccus aminovorans* (*Pa*) was more likely to be present and more abundant in the gut microbiota during *Vc* infection (7). The association between *Pa* and *Vc* is highly unusual because most of the native gut microbiota is typically displaced by secretory diarrhea during cholera (5, 15). To determine the underlying mechanisms driving these correlative clinical findings, we evaluated the relationship between *Pa* and *Vc* in co-culture and determined the effects of *Pa* on *Vc* infection outcomes with *in vivo* models. Here we show that *Pa* interacts directly with *Vc* to form dual-species biofilm structures with previously uncharacterized features. Moreover, *Vc* colonization inside the animal host is enhanced by the presence of *Pa* in the small intestine, and this effect is dependent upon *Vc* biofilm production. Our findings demonstrate a plausible mechanism by which a gut microbe specifically associates with *Vc*, and this reinforces our microbiome analysis in humans that identified *Pa* as highly associated with infected individuals. Our findings also demonstrate that interactions between these two species have the potential to directly impact *Vc* pathogenesis and alter outcomes of *Vc* infection in humans.

## Results

### *P. aminovorans* is differentially abundant in individuals with active *V. cholerae* infection

*Paracoccus* is a genus of soil microbes found in low abundance in the gut microbiome of humans (16, 17). Our previously published analysis of stool gut microbes from household contacts of cholera patients identified *P. aminovorans (Pa*) as an unexpectedly abundant gut microbe during active *V. cholerae* (*Vc*) infection, and this organism was rarely found in uninfected participants (7). In this prior study, we used a support vector machine model with recursive feature elimination to learn patterns of relative abundance of operational taxonomic units (OTUs) that distinguished infected (defined as *Vc* DNA detected in stool, or culture positive) from uninfected persons (*Vc* DNA undetected in stool). The model was trained on a subset of study participants and tested on another subset in a hold-out validation. Here, we have extracted data from this prior study to examine separately the *Pa* OTUs in infected compared to uninfected persons. *Pa* abundance was significantly higher as a proportion of the total sequencing reads in the stool of infected participants (6/22, 27%) of infected household contacts had detectable *Pa* compared to only 5.6% (2/36) of uninfected individuals (Figure 1A). The ratio of *Pa* to *Vc* abundance present during infection was variable and averaged 1:1 (Figure 1B). These findings were particularly interesting because typically there is a drastic reduction of nearly all gut microbes during active *Vc* infection (5, 15) due to secretory diarrhea, oral rehydration solution ingestion, and *Vc* infection itself, and yet here *Pa* was found in an increased abundance in some actively infected participants. Based on these findings, we hypothesized that *Pa* may be resistant to displacement from the gut during infection. While our previous study demonstrates a positive correlation between *Pa* in human stool and *Vc* infection, a causal relationship between this gut microbiota species and *Vc* infection had not been previously established.

**Figure 1.**
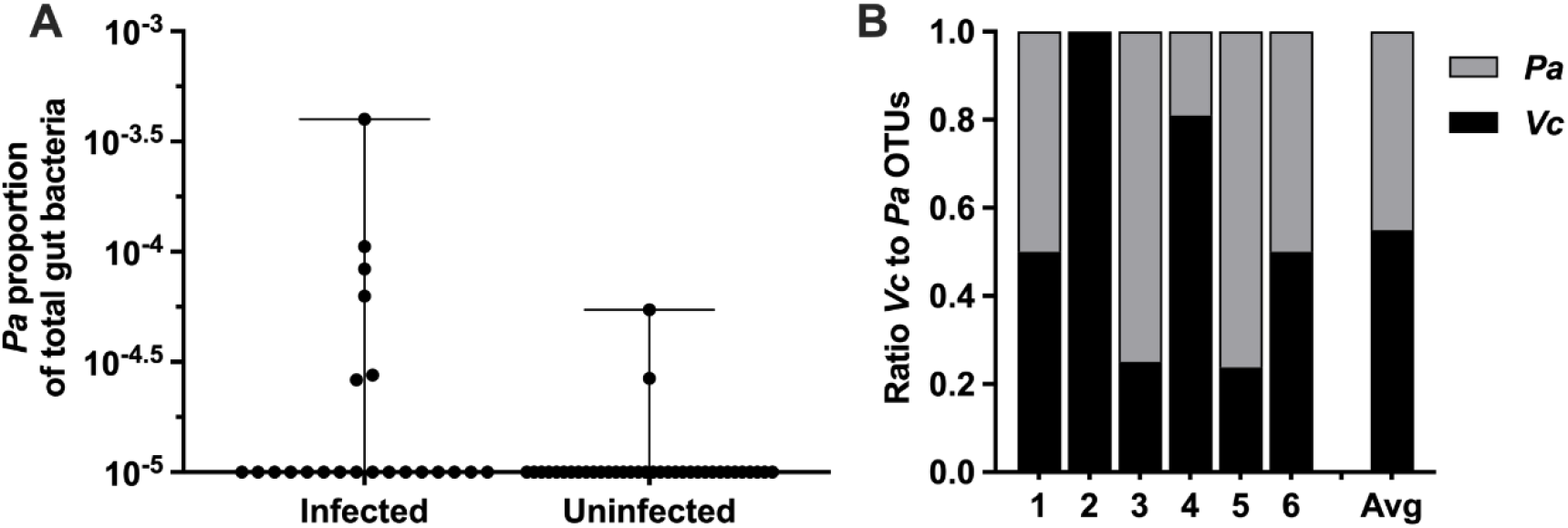
*Pa* is more abundant in persons with *Vc* infection compared to uninfected persons. In a prior study of household contacts of cholera patients in Bangladesh (7), *Pa* was identified as differentially abundant using a support vector machine model with recursive feature elimination in order to discriminate patterns of microbial taxa relative abundance that distinguished infected from uninfected persons. The microbiota was assessed using 16S rRNA in rectal swabs collected from individuals with *Vc* infection (*n* = 22) compared to uninfected individuals (*n* = 36). In this study, total sum normalization was applied to OTU counts from each sample, and a median of 37,958 mapped reads per sample were generated (7). Based on this sequencing data, the estimated limit of detection for a *Pa* OTU is 2.0×10^−5^. **(A)** Relative abundance *Pa* in infected and uninfected individuals and **(B)** ratio of *Vc* to *Pa* in six *Vc*-infected persons. All data points are shown and boxes indicate interquartile range. Bars mark the maximum and minimum values. These values were compared with non-parametric unpaired Mann-Whitney U testing, *P* <0.01.

### *Pa* increases *Vc* host colonization

We modified a well-established infant mouse colonization model (18) to assess whether the presence of *Pa* in the small intestine would promote *Vc* host colonization. First, we isolated a spontaneous streptomycin resistant (Strep^R^) mutant derived from the ATCC type strain of *Pa* for selection and enumeration of *Pa* following host colonization. Infant mice (3-day old) were intragastrically inoculated with *Pa* (10^7^ colony forming units [CFUs]) every 12 hours for 4 doses (0, 12, 24, 36 hours). At 24 and 48 hours after the first inoculation, small intestines from these animals were dissected and homogenized. Gut homogenates were serially diluted and plated on medium containing streptomycin to assess *Pa* colonization. Strep^R^ *Pa* colonies (>10^6^ CFUs/small intestine) were recovered at these two time points (Figure 2A), and no Strep^R^ colonies was detected in the mock treated group, indicating that *Pa* successfully and stably colonized the small intestines of these animals using these methods. Unlike previous studies (9, 10), pretreatment with antibiotics did not change the outcome of *Pa* colonization (not shown). Sequencing analysis of the mouse small intestines demonstrated no significant change in the microbial composition and diversity with and without *Pa* colonization (Supplemental Figure 1 A and B).

**Figure 2.**
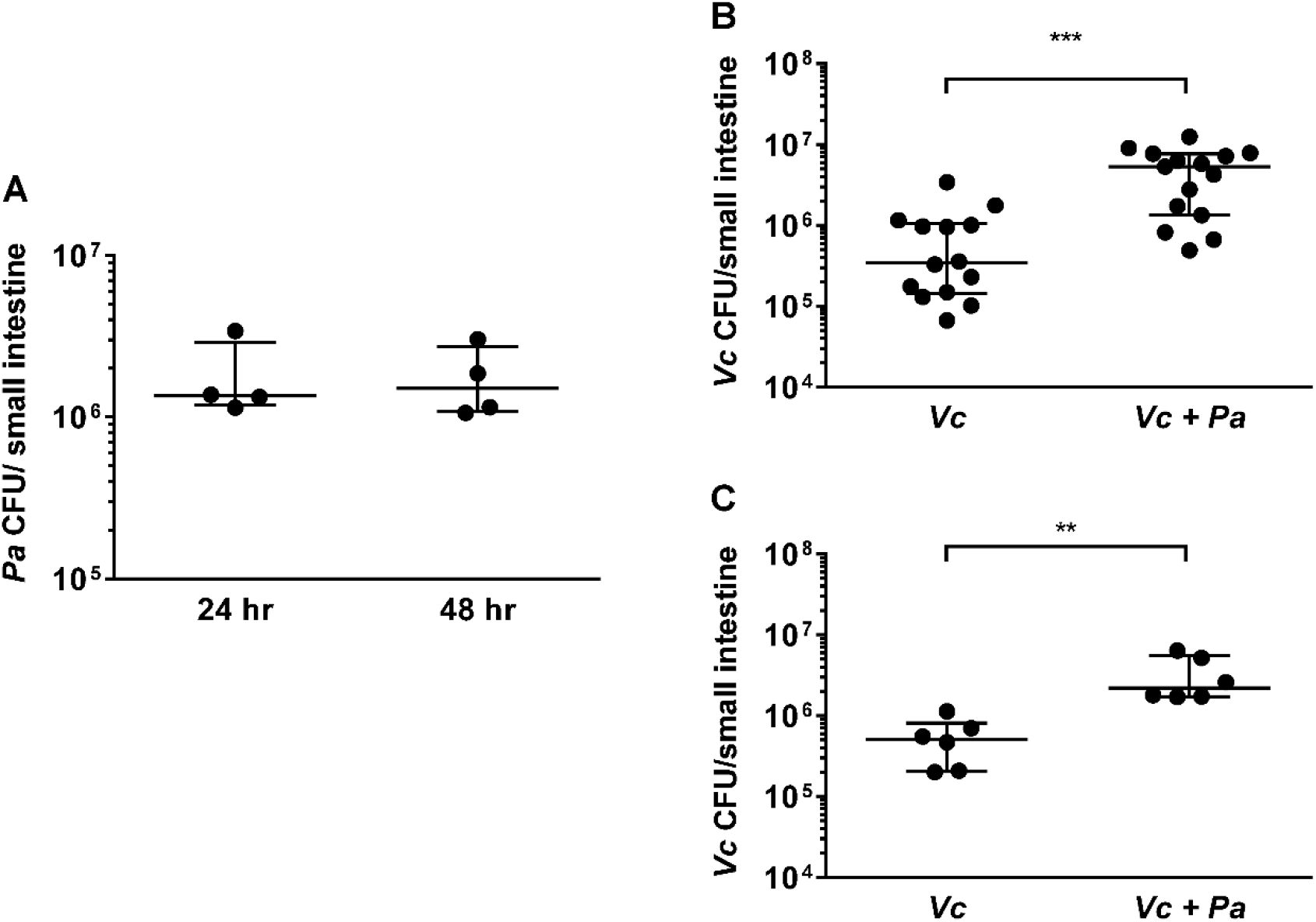
The presence of *Pa* enhances *Vc* colonization in the infant mouse intestine. **(A)** 3 day-old infant mice were intragastrically inoculated with 10^7^ CFU of *Pa* every 12 hours for a period of 24 hr (2 doses) or 48 hr (4 doses). At each time point mice were sacrificed and CFU were enumerated by plating serial dilutions of small intestine samples on selective media. **(B)** 3 day-old infant mice were intragastrically inoculated 4 times with LB or 10^7^ CFU of *Pa* for every 12 hours, and subsequently infected with 10^6^ CFU of WT *Vc*. Mice were sacrificed 20-24 hr post-infection and the small intestine samples were taken to enumerate *Vc*. Bars on graphs depict median value with 95% confidence interval (CI) and individual data points plotted. Unpaired non-parametric *t*-test (Mann-Whitney); *** *P* ≤ 0.0001. (C) *Vc* was inoculated intragastrically into the animals alone or together with *Pa* in a 1:1 ratio. After 24 hours, enumeration of *Vc* was performed as described above. Unpaired non-parametric *t*-test (Mann-Whitney); ** *P* ≤ 0.005.

We then evaluated if pre-colonization by *Pa* would influence *Vc* colonization in the small intestine. *Pa* pre-colonization in the infant mice was established over a 36-hour period as described above. Negative control animals were inoculated with sterile media in place of *Pa* over the same dosing schedule. Twelve hours after the last *Pa* inoculation (i.e., 48 hours after the first *Pa* inoculation), these animals were infected with *Vc* (10^6^ CFU) to evaluate whether pre-colonization with *Pa* had an impact on *Vc* colonization. Although we do not fully understand the exact composition and growth dynamics of *Vc* and *Pa* inside the human gut, the pre-colonization/infection scheme was aimed to closely simulate the ratio of *Pa* to *Vc* observed in the gut microbiota of *Vc* infected humans (Figures 1 and 2A). Comparing *Pa* pre-colonized mice to the control group, there was a significant increase (~10-fold, p≤0.0001) of *Vc* colonization in the mice pre-colonized with *Pa* (Figure 2B) 24 hours after infection. This enhanced intestinal colonization by *Vc* in the *Pa*-colonized mice was observed as early as 6 hours after infection and maintained throughout the colonization period (Supplementary Figure 2).

We reasoned that it was also possible for *Vc* and *Pa* to encounter one another in the environment before entering the host. To model this scenario, *Vc* was mixed with *Pa* in 1:1 ratio, and the mixture was used immediately for animal infection. In agreement with the results obtained with the *Pa* pre-colonization model, *Vc* intestinal colonization was significantly higher when coinfected with *Pa* than without *Pa* (Figure 2C). Given *Pa* colonization did not overtly change the overall composition of the gut microbiota (Supplementary Figure 1), collectively, our results demonstrate that the presence of a single gut microbiota species is sufficient to increase *Vc* host colonization. Our findings also illustrate that our approach to microbiome studies in humans (6, 7) can be used as a predictive tool to identify gut microbes that alter *Vc* virulence.

### *Pa* promotes *Vc* biofilm formation

To investigate if the increased *Vc* intestinal colonization is due to direct interactions between *Vc* and *Pa*, these two species were co-cultured and allowed to propagate for three days where both planktonic growth and pellicle formation (i.e., biofilm formation at the air-liquid interface) of both species was monitored. There was a small difference (< 2-fold) in growth in the planktonic phase of either *Vc* or *Pa* in the co-cultures when compared to the cultures containing a single species (Figure 3A). However, the *Vc/Pa* co-culture formed a pellicle that was visibly thicker and more robust than that formed by *Vc* monoculture (photo shown in Figure 3B). The *Pa* monoculture did not form a visible pellicle. The co-culture pellicle samples were carefully lifted and removed from the culture medium, washed, and agitated to release single cells for enumeration of each species. Compared to *Vc* monoculture, the co-culture samples contained over 50-fold more *Vc* cells while the ratio of *Vc* to *Pa* approached to approximately 1:1 (Figure 3C). Moreover, only a small fraction (0.01%) of *Vc* and *Pa* could be washed off from the isolated pellicles (Figure 3D), suggesting that these species are tightly integrated into the pellicle.

**Figure 3.**
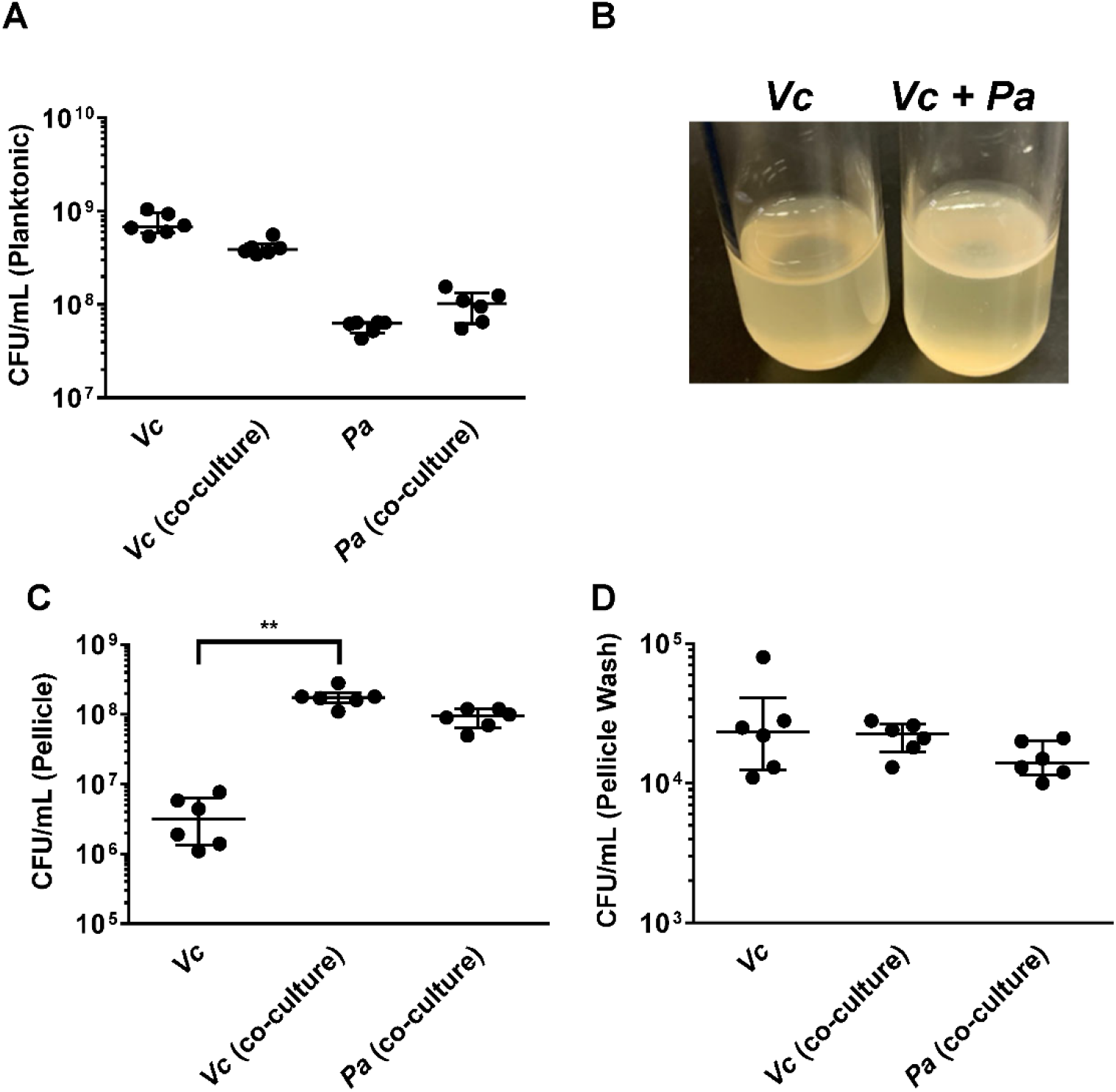
*Pa* promotes biofilm formation of *Vc*. **(A)** Planktonic cell counts from cultures used for pellicle analysis of *Vc* and *Pa* grown together or in monoculture. **(B)** Representative images of pellicles formed by *Vc* grown in monoculture and in co-culture with *Pa*. CFU counts of each strain in **(C)** pellicle samples and **(D)** spent medium used to wash the pellicle. Bars on graphs depict median value with 95% confidence interval (CI) and individual data points plotted. Unpaired non-parametric *t*-test (Mann-Whitney): ** *P* ≤ 0.005.

Based on the above data, we hypothesize that *Vc* and *Pa* form dual-species biofilms at the air-liquid interface. This is unexpected because *Vc* is known to form a clonal community in both *in vitro* and *in vivo* biofilms and these are known to exclude other species including even planktonic *Vc* cells (19, 20). To test this hypothesis, we transferred the co-culture pellicles onto coverslips for imaging with confocal microscopy (Figure 4). All cells in the pellicle were stained with FM 4-64 membrane dye, and *Vc* cells were differentiated from *Pa* using a constitutively produced mNeonGreen reporter (21) expressed from a neutral *Vc* locus (22). In the *Vc/Pa* co-culture pellicles, we observed a continuous film structure spanning the entire pellicle (Figure 4A). Notably, cocci-shaped *Pa* cells were clearly visible in the co-culture pellicle (Figure 4B-C), consistent with the CFU quantification in Figure 3A. Interestingly, *Pa* cells were found throughout the pellicle, with a higher abundance in the bottom layer (Figure 4D-F), and always in close association with *Vc* cells. In summary, we found that *Vc* and *Pa* coexist stably in the pellicle structure and this relationship may explain the mechanism by which *Pa* resists displacement in humans during active *Vc* infection.

**Figure 4.**
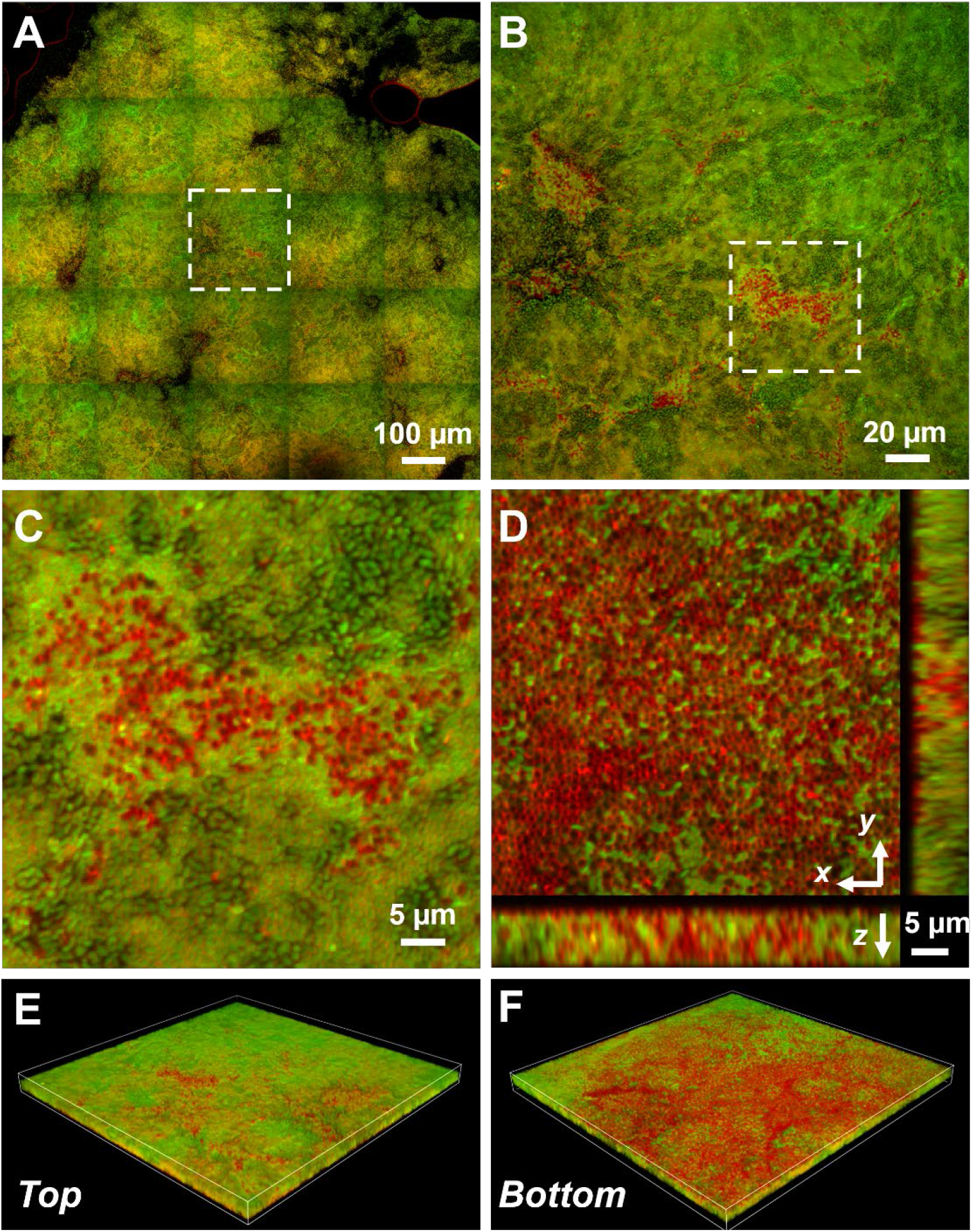
Representative microscopy images of *Vc* and *Pa* dual-species pellicles. **(A)** Large-scale cross-sectional image of the internal structure in a co-culture pellicle. All cells are stained with FM 4-64 and *Vc* cells constitutively express mNeonGreen. Therefore, the red signal in the overlay image corresponds primarily to *Pa* cells. **(B)** Zoom-in view of the region highlighted in A. **(C)** Zoom-in view of the region highlighted in B. **(D)** Cross-sectional views of the region shown in C, at the bottom of the pellicle. *Pa* cells exist mainly at the pellicle-liquid interface, with clusters of *Pa* cells penetrating into the interior of the pellicle. **(E-F)** *Top* (E) and *Bottom* (F) view of the co-culture structure shown in B, rendered in 3D.

Next, we used a standard crystal violet (CV) microtiter plate assay (23) to quantitatively evaluate how *Vc* and *Pa* interact under pellicle forming conditions. *Vc* and *Pa* were simultaneously inoculated into the wells of microplates in two different *Vc*:*Pa* ratios (1:1 and 1:10). We also tested if the viability of *Pa* was crucial for this interaction by using heat-killed *Pa* as a control. Consistent with our pellicle compositional analysis, *Vc* formed a more robust biofilm than *Pa* under these conditions as demonstrated by increased CV staining in wells containing *Vc* only compared to wells containing *Pa* only (Figure 5A). Importantly, CV staining was increased in wells containing *Vc* and live *Pa* compared to wells with *Vc* only, in a concentration-dependent manner (Figure 5A). In contrast, CV staining was not different in wells containing *Vc* and heat-killed *Pa* compared to wells with *Vc* only (Figure 5A).

**Figure 5.**
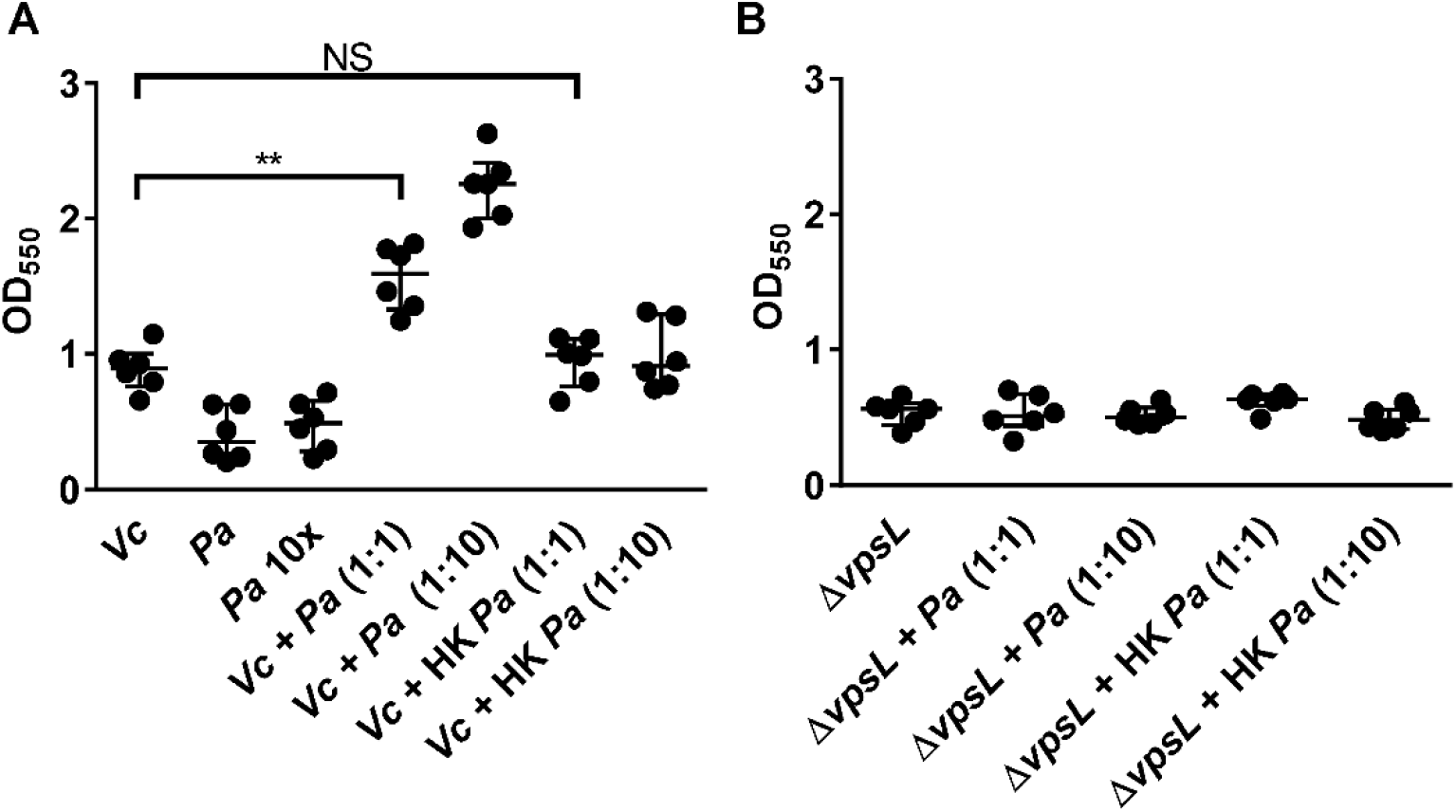
*Pa* increases biofilm production in *Vc*. Crystal violet assays were performed in 96-well microtiter plates to quantify biofilm formation. Overnight-grown **(A)** wild-type *Vc* or **(B)** *ΔvpsL* mutant and *Pa* cultures were diluted to a final concentration of 10^6^ CFU in a total volume of 200 μL/well. In samples containing a 1:10 ratio of *Vc*/*Pa*, *Pa* was diluted to a final concentration of 10^7^ CFU. Samples with heat-killed (HK) *Pa* are specified in the x-axis. Microtiter plates were incubated at 37 °C for 24 hr. Crystal violet staining and ethanol solubilization were performed as previously described (23). Absorbance of the crystal violet stain was measured at 550 nm using a Biotek Synergy HTX plate reader. Data is represented with horizontal lines indicating the mean with standard deviation. Unpaired *t*-test (Mann-Whitney); \*\**P* ≤ 0.005.

To replicate our mouse experiments (Figure 2), we also tested if the order in which the two species encounter one another is critical for the *Vc* biofilm enhancement phenotype. *Pa* was grown in wells 24 hours before the addition of *Vc*. As in our previous results in the co-inoculation experiment, an increase in CV staining was observed in wells in which the two species were added sequentially, but not in the wells with *Vc* only (Supplementary Figure 3). Moreover, wells pre-incubated with heat-killed *Pa* and subsequently inoculated with *Vc* had no increase in CV staining compared to wells inoculated with *Vc* alone (Supplementary Figure 3). Together, our biofilm quantification data suggests that the presence of *Pa*, regardless of the order of encounter, results in an enhanced biofilm formation of *Vc*.

### *Vibrio* exopolysaccharide is essential for a stable biofilm structure formed by *Vc* and *Pa*

To understand what biofilm component is required for the enhancement of biofilm production in *Vc/Pa* co-culture, we repeated the above experiments with a Δ*vpsL Vc* mutant that cannot produce the *Vibrio* exopolysaccharide (VPS) necessary for mature biofilm formation (24). In contrast to what we observed with a *vpsL*^+^ strain, there was no significant increase in CV staining in wells with both *ΔvpsL* mutants and *Pa*, when compared to wells with the *ΔvpsL* mutants only (Figure 5B).

To further investigate the role of VPS in promoting co-culture biofilms, we stained the co-culture pellicle *in situ* with Wheat Germ Agglutinin (WGA), a common stain for VPS (which contains GlcNAc moieties, (25)). To avoid spatial overlap with the membrane stain (excited at 561 nm), the *Vc* cells used in this experiment express a cyan-fluorescent protein SCFP3A cytosolically (excited at 445 nm), and the WGA is conjugated to Oregon Green (excited at 488 nm). Figure 6A shows a 3D view of a large area of a *Vc/Pa* co-culture pellicle with WGA staining. We then compared the intensity of WGA staining in areas with different compositions of *Vc* and *Pa*. Area 1 in Figure 6A represents a location where *Vc* is the predominant species and *Pa* abundance is low (i.e., FM 4-64 staining overlapped entirely with SCFP3A signal), while areas 2 and 3 in Figure 6A represent locations where *Vc* and *Pa* coexist (i.e., contain regions with FM 4-64 staining but no SCFP3A signal). Surprisingly, the WGA signal intensity was elevated in the *Pa*-rich regions rather than in the *Vc*-rich regions (Figure 6B). Zoom-in view of the *Vc*-enriched region shows the characteristic sub-envelope structures around clusters of *Vc* cells (Figure 6C), consistent with the known VPS morphology in submerged *Vc* biofilms (26). In *Pa*-rich regions, the WGA-signal was stronger, and the VPS structures were more compact (Figure 6C). Importantly, the VPS structure we observed enclose both *Pa* and *Vc* cells, providing an intuitive explanation for how *Pa* cells are incorporated into the pellicle. Together, these results suggested that the physical presence of *Pa* in co-culture pellicle augments the production of VPS in *Vc* cells, leading to increased *Vc* biofilm formation; the *Pa* cells, in turn, rely on VPS to be integrated into the 3D structure of the pellicle.

**Figure 6.**
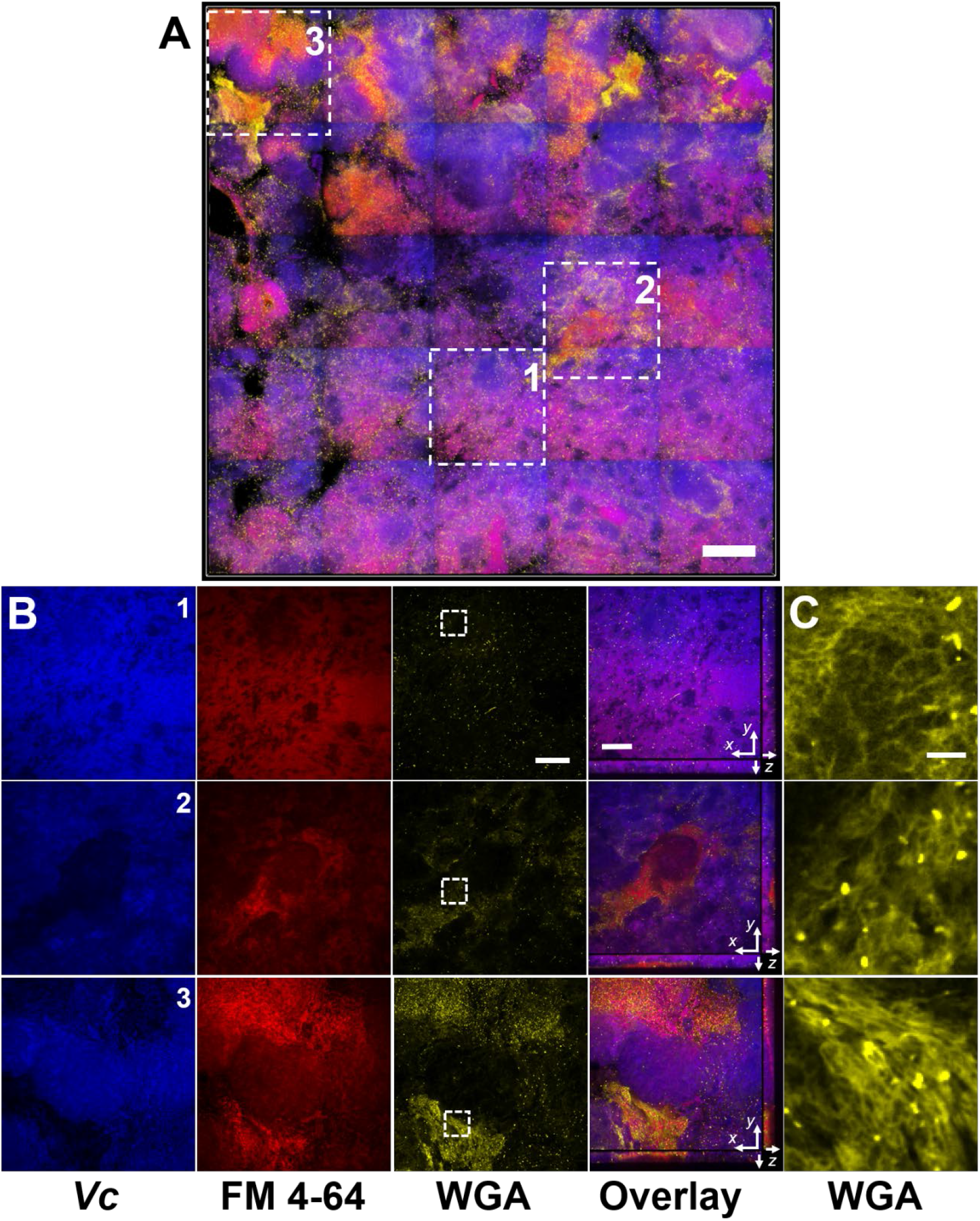
*Vc/Pa* co-culture biofilms depend on VPS. **(A)** Representative top view of a co-culture pellicle with WGA staining, rendered in 3D. *Vc* cells constitutively express SCFP3A cytosolically; all cells were stained with FM 4-64 membrane stain; WGA is conjugated to Oregon Green and shown in yellow. Note that WGA also stained dead cells with an exposed peptidoglycan layer, corresponding to the bright spots in the image. Scale bar: 100 μm. **(B)** Zoom-in view of the three regions 1-3 indicated by the white boxes in A. Shown from *left* to *right* are *Vc* cell fluorescence (SCFP3A), membrane staining (FM 4-64), WGA staining (Oregon Green), and the overlay of the three channels. The overlaid images additionally show the cross-sectional view in the *xz* and *yz* planes. Region 1 contains primarily *Vc* cells, and regions 2 and 3 contain *Pa*-enriched regions. *Pa*-enriched regions demonstrate elevated WGA signal. Intensities in each channel were kept consistent through region 1-3 for comparison. Scale bars: 20 μm. **(C)** Zoom-in view of the highlighted regions in B (white boxes, WGA channel only). Intensities are adjusted to similar level for visualization of the internal structure. Scale bar: 5 μm.

### Enhancement of *Vc* host colonization by *Pa* depends on biofilm exopolysaccharide

Biofilm-grown *Vc* cells are known to be more infectious in humans due to increased resistance to gastric pH and higher expression of virulence factors (e.g., such as the toxin co-regulated pilus, which mediates host colonization) compared to planktonically grown cells (27–29). We hypothesize that because *Vc* biofilm formation is enhanced in the presence of *Pa*, this results in increased virulence inside the host, in a VPS-dependent manner.

To test our hypothesis and measure if the effect of the *Vc/Pa* biofilm interaction impacts host colonization, we compared the colonization efficiency between wild-type (WT) or the *ΔvpsL* mutants in infant mice with and without *Pa* pre-colonization. As shown previously (30), the *ΔvpsL* mutant was able to colonize the mouse small intestine equally as well as the WT *vpsL*^+^ strain, confirming that the VPS is not absolutely required for host colonization when *Vc* was administered to the animals alone. In contrast, while *Pa* increased WT *vpsL*^+^ *Vc* colonization, the *ΔvpsL* mutant did not exhibit the enhanced colonization phenotype in the *Pa* pre-colonized mice (Figure 7A). Similar results were observed using the coinfection model; when *Vc ΔvpsL* mutants were coinfected with *Pa*, there was no increase in host colonization (Figure 7B). Together, we concluded that the enhancement of *Vc* intestinal colonization in the presence *Pa* is dependent on the *Vibrio* exopolysaccharide, in line with our *in vitro* data.

**Figure 7.**
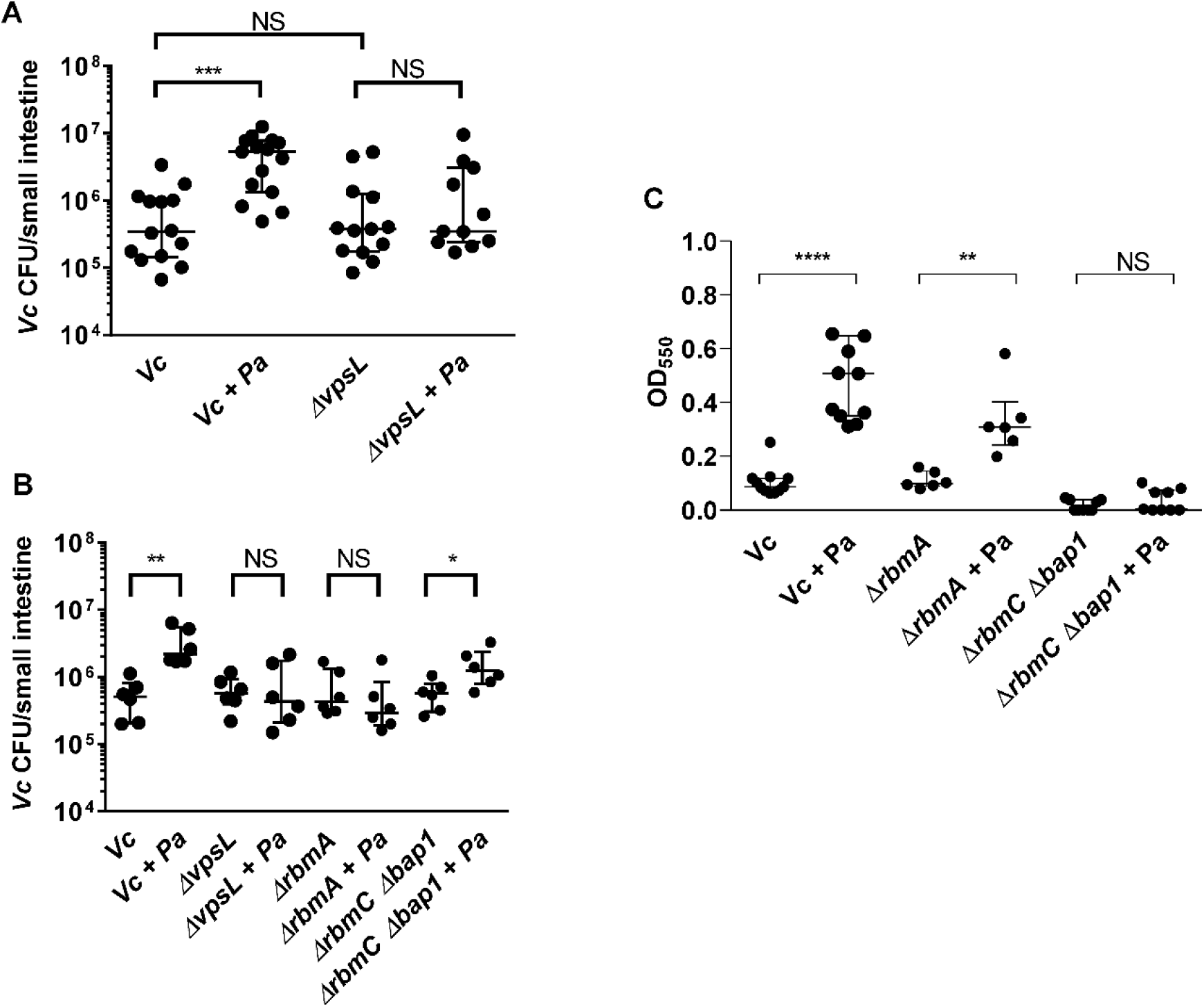
Enhanced *Vc* intestinal colonization in the presence of *Pa* is dependent on VPS and accessory matrix proteins. **(A)** 3 day-old infant mice were intragastrically inoculated with LB or 10^7^ CFU of *Pa* every 12 hours for a period of 48 hours, and subsequently infected with 10^6^ CFU of a *Vc* strain defective for extracellular matrix production (*ΔvpsL*). Mice were sacrificed 20-24 hr post-infection and small intestine samples were taken to enumerate *Vc* CFU. Data from infection with the wild-type *Vc* strain (Figure 2B) is shown again here for comparison purposes. **(B)** *Vc* wild-type (WT) or different biofilm mutants were mixed with *Pa* in 1:1 ratio, and the mixture was used immediately for animal infection. Mice were sacrificed 20-24 hr post-infection and small intestine samples were taken to enumerate *Vc* CFU. Each symbol represents an individual mouse and data is represented with horizontal lines indicating the median with a 95% confidence interval. Unpaired non-parametric *t*-test (Mann-Whitney); ns *P* > 0.05, *** *P* ≤ 0.001. Data from infection with the wild-type *Vc* strain (Figure 2C) is shown again here for comparison purposes. **(C)** Crystal violet assays performed in 96-well plates to quantify pellicle formation. Overnight cultures of *Vc rbmA-, rbmC-bap1*- and *Pa* were diluted in fresh LB and plated as 200 μL/well. Samples were co-cultured in either 1:10 ratios of Vc/Pa and incubated at 37 °C for 24 hr. Crystal violet staining was then performed and absorbance of the stain was measured at 570 nm. Horizontal lines indicating mean with standard deviation are shown. Unpaired *t*-test; \*\*\*\**P* ≤ 0.0001, \*\**P* ≤ 0.05.

### Accessary biofilm matrix proteins are involved in *Pa* and *Vc* interaction

Mature *Vc* biofilm is stabilized with a variety of accessary matrix proteins in addition to the VPS (26, 31, 32). To interrogate the roles of these components in the interactions between *Vc* and *Pa*, we tested mutants lacking the cell-cell adhesion proteins RbmA (26, 31, 33) and mutants lacking surface adhesion redundantly conferred by RbmC and Bap1 (26, 32, 33) for their ability to increase biofilm formation in the presence of *Pa* using CV assays (Figure 7C). When compared to the wells containing the Δ*rbmA* mutant alone, the CV staining was higher in the wells with both the Δ*rbmA* mutant and *Pa*. However, the increase was not as high in the Δ*rbmA* mutant when compared to that in WT *Vc*/*Pa* co-culture (Figure 7C). Furthermore, the presence of *Pa* did not increase CV staining in the wells containing the Δ*rbmC* Δ*bap1* mutants (Figure 7C).

We then performed the infant mouse colonization experiments with *Vc* biofilm matrix protein mutants using the *Pa* coinfection model to test the roles of these proteins *in vivo*. For this series of experiments, each *Vc* biofilm mutant was coinfected into the animals with *Pa* in 1:1 ratio. In agreement with our *in vitro* results, while host colonization was significantly higher for WT *Vc* coinfected with *Pa* than without *Pa* (Figure 2C), Δ*rbmA* mutants did not show any increase in host colonization when coinfected with *Pa*, and Δ*rbmC* Δ*bap1* mutants demonstrated reduced enhancement in colonization during coinfection with *Pa* (Figure 7B). These results indicate that the ability of *Vc* to form a structurally intact biofilm is important for the enhancement of colonization facilitated by the presence of *Pa*.

## Discussion

Evidence that the composition of the gut microbiota influences the clinical outcomes of enteric infections in humans is accumulating (34, 35). Several studies have identified commensal species and underlying colonization resistance mechanisms that could be protective against *V. cholerae* (*Vc*) infection. While these studies suggest that microbiota species reduce *Vc* virulence through various mechanisms during the early stages of infection (4, 5, 14), the precise role of these colonization resistance mechanisms in impacting susceptibility to cholera in humans has only begun to be appreciated. For instance, a bacterium in the genus *Blautia* was recently found to encode functions that confer colonization resistance (e.g., bile salt hydrolase) to *Vc* infection (4). Consistent with this finding, our previous stool microbiome study has also independently identified that one of the species in the genus *Blautia* is correlated with decreased susceptibility to *Vc* infection (6, 7).

While previous studies have identified microbiota-associated mechanisms that are protective against *Vc* infection, examples of interactions between *Vc* and a human-associated microbiota species that increases *Vc* pathogenicity are scarce. Although *Escherichia coli* and *Vc* are believed to reside in different intestinal niches, one previous study showed that an atypical *E. coli* isolated from a mouse that does not ferment lactose can increase the virulence of a quorum-sensing (QS) defective *Vc* strain N16961 (36). How QS-proficient *Vc* strains, which are prevalent in toxigenic clinical isolates (37), respond to typical *E. coli* in the human gut remains to be studied. In contrast, a recent study showed that *E. coli* motility facilitates aggregation of these two organisms in a dual-species biofilm, but there was no impact of such aggregation on *Vc* intestinal colonization (38). Indeed, coaggregation between *Vc* and other microbiota species has been observed (39), but these associations are not known to have a direct influence on *Vc* pathogenicity. This is consistent with our prior human studies in which *E. coli* species were present in the gut microbiota of persons during active *Vc* infection, but these were not correlated with active *Vc* infection (7). Our findings highlight the importance of coupling mechanistic studies (*in vitro* and animal models) with human microbiome data analysis to pinpoint the relevant species and interactions involved in enteric infections.

Here we show that the presence of a human gut microbe *Pa* promotes *Vc* host colonization, which is consistent with our prior human study in which *Pa* was more likely to be present in persons infected with *Vc*. This raises the possibility that uncharacterized interactions between *Vc* and members of the gut microbiota may exacerbate *Vc* virulence and contribute to increased morbidity. Our current study also establishes a plausible mechanism used by *Pa*, and perhaps other gut microbes, to increase the virulence of *Vc* through induction of biofilm formation, a physiological state in which *Vc* is known to increase expression of other virulence factors critical for human infection and disease (27, 28). *Vc* biofilms have also been demonstrated to deform and even damage tissue-engineered soft epithelia mimicking the host tissue (40), suggesting that *in vivo*-formed biofilm structures could negatively impact host gut physiology.

While VPS and other biofilm components are not usually considered critical host colonization factors, we found that these macromolecular structures were essential for the enhancement of *Vc* host colonization induced by *Pa*. Whether these components mediate other *Vc*-gut microbe interaction has not been studied. Interestingly, many gut microbes appear to predominantly exist in the form of mixed-species biofilms on mucosal surfaces (41), suggesting microbiota-induced biofilm enhancement could play a major role in modulating virulence of other pathogens. Many structural components, regulatory factors, and signaling transduction pathways that control biofilm formation in *Vc* have been well characterized (42), and these factors could be targeted for manipulation by other gut microbes that modulate *Vc* virulence. For example, 3,5-dimethylpyrazin-2-ol (DPO) was recently discovered as a new class of *Vc* quorum-sensing autoinducer that binds to the transcription factor VqmA to activate expression of *vqmR*, which encodes a small regulatory RNA that downregulates *Vc* biofilm formation. The VqmA-VqmR system can be activated both *in vitro* by *E. coli* and *in vivo* by *B. obeum* (5, 43), and results in suppression of biofilm formation. Interestingly, *Pa* demonstrates the opposite tendency by promoting *Vc* biofilm formation, with implications for the enhancement of *Vc* colonization, in contrast to other commensal bacteria.

Many aspects of the *Vc*/*Pa* interaction are still unclear. What is the selective advantage that fosters the formation of dual-species biofilm? Investigation of the structure-function relationship in other multispecies biofilms, such as dental biofilms, demonstrates a coordinated organization of each species that allows for optimal nutrient and oxygen usage, as well as mechanical stability (44, 45). While we did not observe any growth yield enhancement in the planktonic phase of the co-culture, there was a significant increase of *Vc* and *Pa* abundance in the co-culture pellicle at the air-liquid interface. Thus, a possible driving force of this interaction could be the optimization of nutrient sharing and distribution, or removal of toxic metabolites accumulated during growth. The exact mechanism used by these two species to detect and coordinate with each other remains unclear. Secreted small molecules produced by *Pa* do not appear to impact *Vc* as evidenced by our prior studies in which *Vc* cultured in *Pa* spent-cell supernatant did not yield result in increased biofilm formation (7). Therefore, we surmised that the close physical association between *Vc* and *Pa* cells in space in the co-culture pellicles is required for the enhanced biofilm formation. This hypothesis is supported by our microscopy analysis. The *Vc/Pa* interaction has two reciprocal aspects: First, *Pa* activates the production of VPS in *Vc* cells, leading to enhanced pellicle formation. Future characterizations of *Pa* could potentially elucidate the underlying molecular mechanism of this effect *Pa* has on *Vc*. Second, in order to be integrated into the pellicle structure, *Pa* cells seem to physically interact with VPS. Interestingly, we have shown that VPS staining signals are stronger in the *Pa*-rich regions than in the *Vc*-rich regions. This could be explained either by a stronger attraction between VPS and *Pa* than between VPS and *Vc* cells, or by activation of VPS biosynthesis genes in *Vc* cells in the vicinity of the *Pa* cells, or both. Future biochemical and biophysical studies to investigate this relationship may provide new insights about the interactions between *Pa* and *Vc* biofilm, and about pathogen-gut microbe interactions in general. Other members of the *Paracoccus* genus are known to form biofilms and encode adhesins to facilitate surface attachment (46, 47), and the potential role of these adhesins in facilitating interaction with *Vc* remains to be studied.

In conclusion, we describe a novel interaction between *Vc* and a gut microbe found in high abundance in *Vc*-infected persons that leads to a significant change in *Vc* biofilm behaviors, as well as an increase in the virulence of the pathogen. Our findings are also consistent with other observations that rare gut microbial species can have significant impacts on microbial ecosystems (48). This study adds to the growing number of pathogen-gut microbial species interactions that may impact outcomes in human diseases.

## Materials and Methods

### Prior published study sample collection and analysis

In a prior study, we enrolled household contacts of persons hospitalized with cholera at the International Centre for Diarrheal Disease Research, Bangladesh (icddr,b)(7). Briefly, in this previously published study, household contacts were followed prospectively with rectal swab sampling, 30 days of clinical symptom report, and vibriocidal titer measurements, and 16S rRNA sequencing was performed on rectal swab sampling from the day of enrollment in the study (7). Persons with evidence of *V. cholerae* (*Vc*) infection at the time of enrollment in the study were compared to those who did not have evidence of infection in a model to detect gut microbes that were differentially abundant during *Vc* infection (7). *Vc* infection was defined as *Vc* DNA identified on 16S rRNA sequencing or a positive *Vc* stool culture. In this previously published study, we used a machine learning method called a support vector machine (SVM), which utilizes patterns of OTU relative abundance to detect OTUs associated with infected compared to uninfected persons. This SVM was used with a recursive feature elimination algorithm that simplifies models and increases accuracy of the identification of differentially associated OTUs by removing uninformative bacterial taxa (7). For the present study, we re-examined the microbiome data from household contacts at the time of enrollment to quantify the abundance of 16s rRNA reads that mapped to *Pa* OTUs between uninfected study participants and infected participants.

### Strains and culture conditions

All *V. cholerae* (*Vc*) strains used in this study are streptomycin-resistant derivatives of C6706, a 1991 El Tor O1 clinical isolate from Peru (49). The in-frame *ΔvpsL* deletion mutants used in various assays were previously described (50). The *ΔrbmA* and *ΔrbmC Δbap1* mutants were constructed by allelic exchange (51) using specific suicide vectors described before (20, 52). *Vc* strains used for microscopy experiments, Δ*vc1807*∷*P_tac_–mNeonGreen* and Δ*vc1807*∷*P_tac_ –SCFP3A-spec^R^*, were constructed using natural transformation as previous described (22). The *P. aminovorans* (*Pa*) used in our experiments is a Strep^R^ isolate derived from the ATCC type strain (ATCC #49632). *Vc* and *Pa* overnight cultures were grown with aeration in LB at 30°C. Heat-killed strains were incubated at 60°C for 2 hours prior to experimentation. Unless specified, media was supplemented with streptomycin (Sm, 100 μg/ml) and chloramphenicol (Cm, 10 μg/ml) when appropriate.

### Animal studies

For establishing colonization of the microbiota species, 3-day old suckling CD-1 mice (Charles River Laboratories) were fasted for 1 hour, then orally dosed with *Pa* at a concentration of 10^7^ CFU using 30-gauge plastic tubing, after which the animals were placed with a lactating dam for 10-12 hrs and monitored in accordance with the regulations of Tufts Comparative Medicine Services. This inoculation scheme was followed an additional 3 times, for a total of 4 inoculations of *Pa* over the course of 48hrs. After 48hrs, mice were infected with 10^6^ CFU of *Vc*, WT C6706 or mutant strain, or LB as a vehicle control in a gavage volume of 50 μl to evaluate the effect of *Pa* pre-colonization on *Vc* host colonization. At 18-24hrs post-infection, animals were sacrificed, and small intestine tissue samples were collected and homogenized for CFU enumeration. WT *Vc* is *lac*^+^ and appears blue on medium containing X-gal while *Pa* appears white on the same medium. For coinfection experiments, cultures of *Vc* and *Pa* strains were mixed in a 1:1 ratio and mice were orally dosed with a final bacterial count of 10^6^ CFU. Mice were sacrificed 20-24 hours post-infection and small intestine samples were processed as outlined above to evaluate the colonization efficiency of both species.

### Ethics Statement

All animal experiments were performed at and in accordance with the rules of the Tufts Comparative Medicine Services (CMS), following the guidelines of the American Veterinary Medical Association (AVMA) as well as the Guide for the Care and Use of Laboratory Animals of the National Institutes of Health. All procedures were performed with approval of the Tufts University CMS (Protocol# B 2018-99). Euthanasia was performed in accordance with guidelines provided by the AVMA and was approved by the Tufts CMS. The previously published study from which Figure 1 is derived (7) received approval from the Ethical Review Committee at the icddr,b and the institutional review boards of Massachusetts General Hospital and the Duke University Health System. Participants or their guardians provided written informed consent.

### Pellicle composition analysis

To assess pellicle composition, overnight cultures of *Vc* and *Pa* were inoculated into glass culture tubes (18 × 150 mm) containing 2mL LB media in a ratio of 1:10 *Vc* (10^6^) to *Pa* (10^7^) CFU, and co-cultures were allowed to grow statically at room temperature for 3 days. Following static growth, floating pellicles were carefully transferred into sterile 1.5mL Eppendorf tubes containing 1mL LB, and samples were gently spun down to wash away any planktonic bacteria. Planktonic cells were removed, and cell pellets of pellicle samples were resuspended in 1mL of fresh LB media. All samples including supernatant from the pellicle wash step, were serial diluted and plated on Sm/X-Gal media to differentiate *Vc* (blue) and *Pa* (white) colonies.

### Crystal violet biomass assays

Crystal violet biofilm assays were performed as described previously in 96-well flat bottom clear, tissue-culture treated polystyrene microplates (ThermoFisher) (23). In each well, *Vc* (10^6^ CFUs) and/or *Pa* (10^6^ or 10^7^ CFUs) were inoculated into 200μL of medium. For experiments involving spent culture supernatants, *Vc* (10^6^ CFUs) were inoculated into each well containing 200 μL of reconditioned supernatants (80% (v/v) filtered spent culture medium and 20% (v/v) 5× LB). Plates were then sealed using a gas permeable sealing film (BrandTech) and incubated at 37°C. Planktonic culture was removed after 24 hours of incubation, plates were washed with distilled water once. Attached biofilms were stained with 0.1% crystal violet at room temperature for 15-20 min. The amount of biomass adhered to the sides of each well was quantified by dissolving the crystal violet in 95% ethanol and the absorbance of the resulting solution was measured at 550 nm or 570 nm using a plate reader.

### Microscopy

Liquid LB culture of *Vc*, *Pa*, and co-cultures (*Pa*:*Vc* = 10:1) were prepared according to procedures described above. To image pellicles, we used a modified literature procedure (53). Monocultures and co-culture pellicles were first prepared following the procedure described above, except that 3 mL of the culture was incubated in a 5 mL culture tube. After 3 days of incubation at room temperature, the pellicles were carefully picked up by the large end of a 200 μL pipette tip, transferred to a coverslip (22 × 60 mm, No. 1.5), and immediately covered with another square coverslip to prevent drying. The LB medium contained 4 μg/mL FM 4-64 stain (ThermoFisher) to stain all cells. To stain VPS, the LB medium additionally contained 4 μg/mL of Wheat Germ Agglutinin conjugated to Oregon Green (ThermoFisher). The stained biofilms were imaged with a Nikon-W1 confocal microscope using 60× water objective (numerical aperture = 1.20). The imaging window was 221 × 221 μm^2^. For large-scale view, a 5×5 tiling was performed. For zoom-in view, the *z*-step size was 0.5 μm and the pixel size was 108 nm. For large-scale view, the *z*-step size was 1 μm and the pixel size was 216 nm. The mNeonGreen (or SCFP3A) expressed by *Vc* was imaged at 488 nm (or 445 nm) excitation, FM 4-64 at 561 nm, and WGA-Oregon Green at 488 nm with the corresponding filters. All presented images are raw images processed from Nikon Element software.

### Statistics

Error bars in the figures depict the median with a 95% confidence interval as indicted. Based on the experimental design, either standard *t*-test or Mann-Whitney test were used to compare treatment groups as indicated in each figure legend.

## Acknowledgments

We thank members in the Ng and Weil Labs for helpful discussions. We acknowledge Ed Ryan for his assistance in reviewing the manuscript. A.A.W and W-L.N. received support from a Rozan Award from Tufts University School of Medicine for this project. A.A.W. support was provided by AI123494 from the National Institute of Allergy and Infectious Diseases (NIAID). W-L.N was supported by AI121337. J.Y. holds a Career Award at the Scientific Interface from the Burroughs Wellcome Fund.

## Supplemental Figures

**Supplemental Figure 1.**
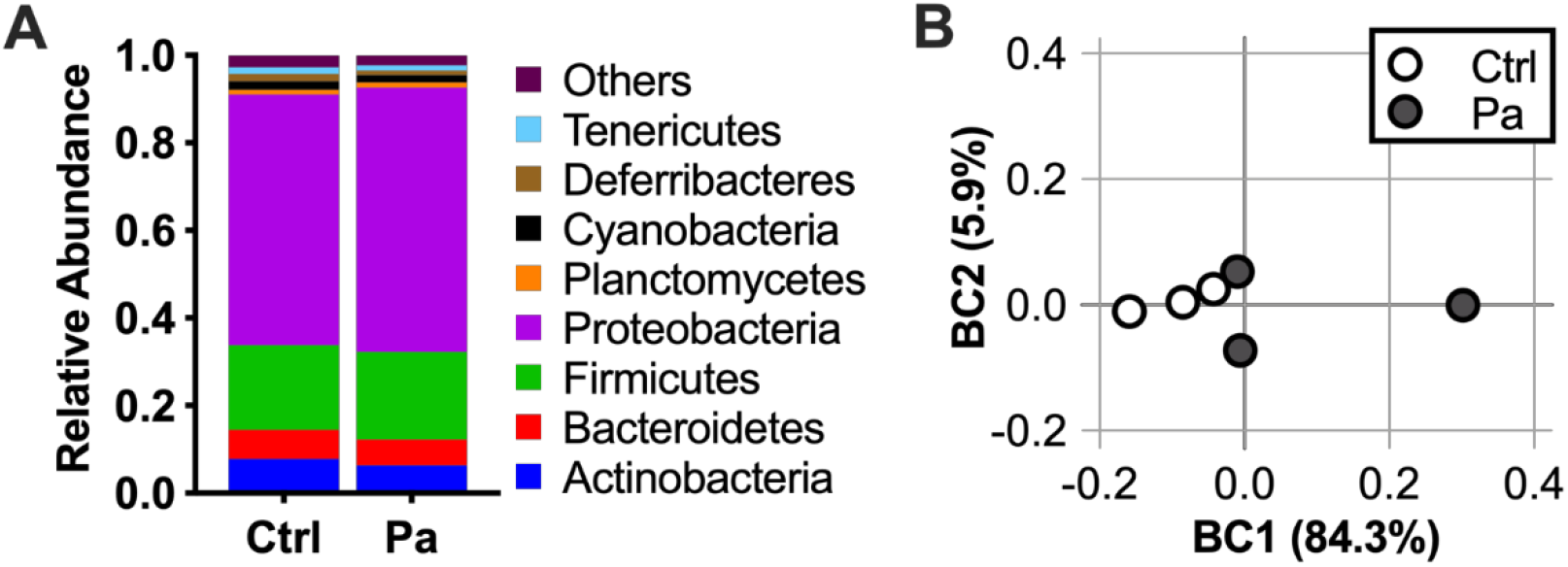
*P. aminovorans* colonization does not significantly alter the mouse gut microbial diversity. Three day-old infant mice were intragastrically inoculated with 10^7^ CFU of *Pa* or sterile LB medium as control (Ctrl) for a period of 12 hours to mimic the length of time between the final *Pa* inoculation and *Vc* infection in mouse experiments. Small intestines were removed, homogenized, and DNA was extracted and sequenced using shallow shotgun sequencing. **(A)** Relative abundance of phylum-level microbes in the small intestines of mice and **(B)** principal component of analysis using Bray Curtis Dissimilarity. N=3 per group. *Pa* is in the Proteobacteria phylum. Two-way ANOVA testing of phylum level-abundance was performed, and all comparisons are not significantly different (P>0.05).

**Supplemental Figure 2.**
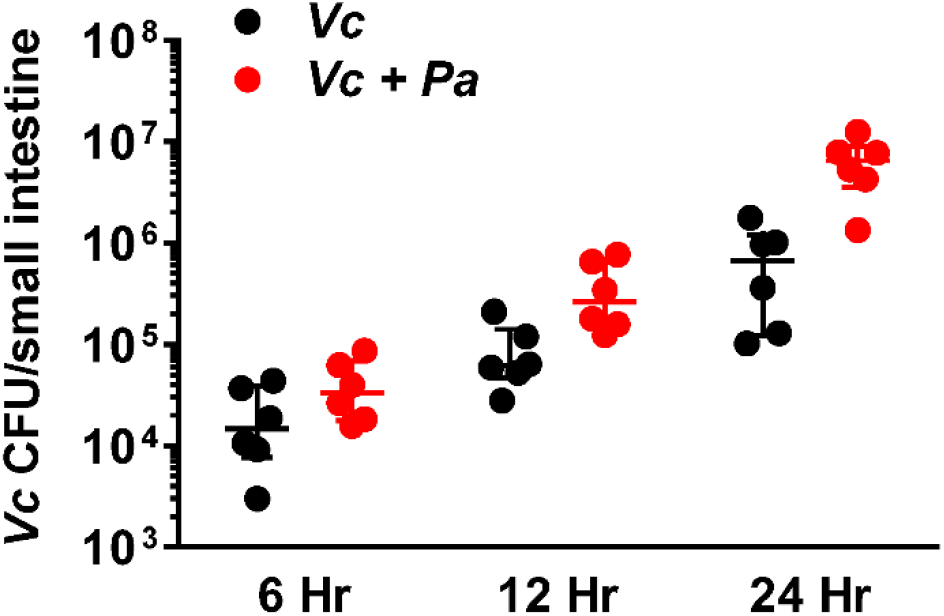
*P. aminovorans* (*Pa*) colonization significantly increases *V. cholerae* (*Vc*) small intestine colonization. Three day-old infant mice were intragastrically inoculated with 10^7^ CFU of *Pa* or sterile LB medium as control for a period of 36 hours. These animals were then infected with *Vc* 48 hours after the first inoculation. At different time points, small intestines were removed, homogenized, and *Vc* were enumerated. Student t-tests were used to analyze these samples. P values are 0.14, 0.035, and 0.004 at 6, 12, and 24 hours respectively (Unpaired non-parametric *t*-test).

**Supplemental Figure 3.**
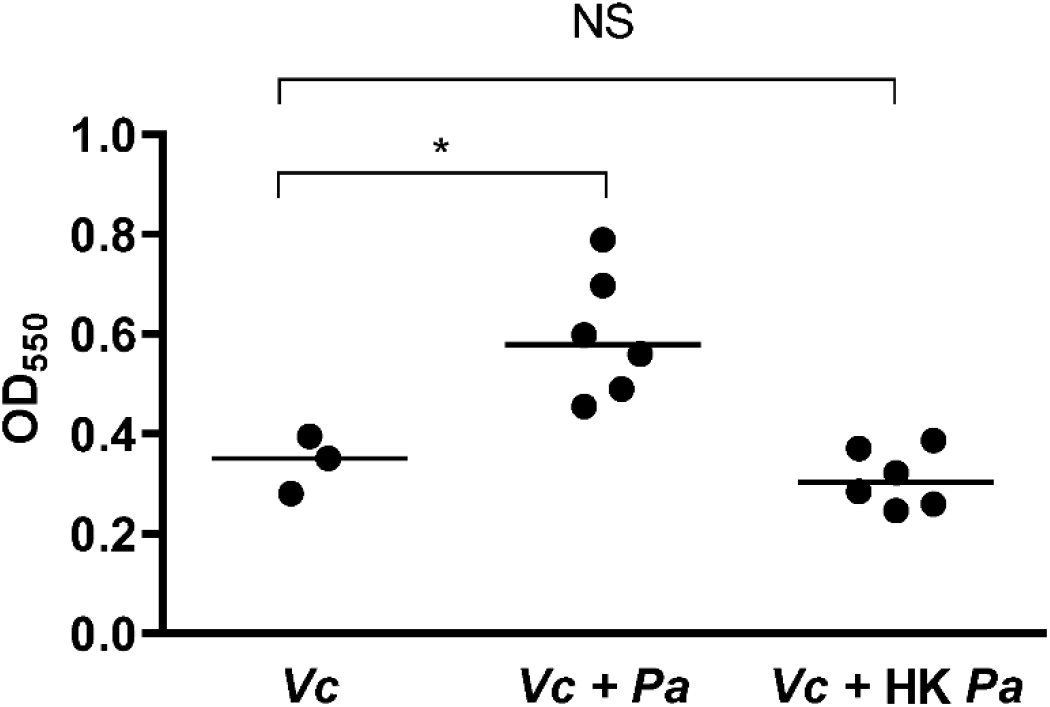
Established *Pa* cultures increase biofilm production in *Vc*. Biofilm formation assays were performed in 96-well microtiter plates. *Pa* was diluted in LB to a concentration of 10^7^ CFU and grown for 24hr before the addition of WT *Vc*. WT *Vc* was then diluted in LB to a concentration of 10^6^ CFU and added to wells. Plates were incubated for an additional 24hr before crystal violet staining to quantify biofilm biomass. Crystal violet staining and ethanol solubilization were performed as previously described (23). Absorbance of the crystal violet stain was measured at 550 nm using a Biotek Synergy HTX plate reader. Samples with heat-killed (HK) *Pa* are delineated by hatched bars. Data is represented with horizontal lines indicating the median with 95% confidence interval. Unpaired non-parametric t-test (Mann-Whitney); * *P* ≤ 0.05.

